# Sympathetic Transmitters Controlling Thermogenic Efficacy of Brown Adipocytes by Modulating Mitochondrial Complex V

**DOI:** 10.1101/161968

**Authors:** Tao-Rong Xie, Chun-Feng Liu, Jian-Sheng Kang

**Author notes:** These authors contributed equally to this work. Correspondence should be addressed to J.-S. Kang, Institute for Nutritional Sciences, Shanghai Institutes for Biological Sciences, Chinese Academy of Sciences, Building 34, Room 315, 294 Taiyuan Road, Shanghai 200031, China.

## Abstract

Obesity is epidemic worldwide as the results of excessive energy intake or inefficient energy expenditure. It is promising to utilize the thermogenic function of brown adipose tissue for obesity intervention. However, the mechanisms controlling the efficacy of norepinephrine-induced thermogenesis in brown adipocytes remain elusive. Here, we demonstrate that the proton ATPase activity of mitochondrial complex V is a key factor, which antagonizes proton leak by UCP1 and determines the efficacy of norepinephrine-induced thermogenesis in brown adipocytes. Furthermore, to avoid unnecessary and undesired heat production, we reveal that ATP as a sympathetic cotransmitter is necessary for the high efficacy and specificity of norepinephrine-induced thermogenesis in brown adipocytes by upregulating the ATP synthase activity of complex V. Thus, we demonstrate the modulation mechanism of thermogenic efficacy in brown adipocytes. These findings imply new strategies for efficiently utilizing brown adipocytes thermogenic capacity, and therapeutic targets for the treatments of obesity and diabetes.

**Highlights:** 1. Norepinephrine (NE) induces heterogeneous responses in brown adipocytes

2. NE activates the H^+^-ATPase activity of mitochondrial complex V

3. Mitochondrial complex V is a key factor in NE-induced thermogenic efficacy

4. ATP as a sympathetic cotransmitter enhances the NE-induced thermogenic efficacy

## INTRODUCTION

Obesity is epidemic worldwide as the results of excessive energy intake or inefficient energy expenditure. Brown adipose tissue (BAT) is the major tissue for cold-induced thermogenesis (energy dissipation as heat) without shivering(1)’(2). Identification of functional BAT in adult humans(3)’(4)’(5) suggests that it is promising to utilize BAT thermogenic capacity for obesity and diabetes treatments(6)’(7).

Currently, the mechanism of thermogenesis in BAT and brown adipocytes (BA) is known in general(2). Cutaneous thermosensory signals for cold stimulation evoke sympathetic nerve firing in BAT via hypothalamus(8). Released neurotransmitter norepinephrine (NE) binds adrenoreceptors of BA, upregulates hormone-sensitive lipase (HSL)-mediated lipolysis via phosphorylation by protein kinase A (PKA) signaling, mobilizes free fatty acids to activate mitochondrial uncoupling protein-1 (UCP1), and converts electrochemical potential energy stored in mitochondrial proton gradient to heat(2)’(9). However, our previous study demonstrates a low thermogenic efficacy of NE-induced thermogenesis in BA (10), which may partially account for the failures of trials to utilize BAT thermogenic capacity for obesity intervention(11),(12). Therefore, the regulatory mechanism of thermogenesis in BA is still largely unknown. In this study, we demonstrate that proton-ATPase function of mitochondrial complex V accounts for NE-induced heterogeneous changes in BA and that sympathetic cotransmitter ATP enhances the efficacy of thermogenesis in BA.

## RESULTS

### NE induces heterogeneous responeses in BA

We consistently observe that NE induces heterogeneous changes of mitochondrial membrane potentials (MMP) in BA (Figure 1), which is monitored with thermoneutral rhodamine 800 (Rh800, a thermal-insensitive mitochondrial marker and MMP sensor)(17) (10). Obviously, there are two MMP populations (depolarization and hyperpolarization) of NE-induced thermogenesis in BA (Figure 1A-H), in line with previous studies (18) (10). The subpopulation of MMP depolarization can be explained by the activation of UCP1 dissipating electrochemical potential energy of proton as heat. The diverse extent of MMP depolarization evoked by NE may be explained by heterogeneity and different NE affinity of β adrenergic receptors(19) or the heterogeneous levels of UCP1 expression(20).

**Figure 1.**
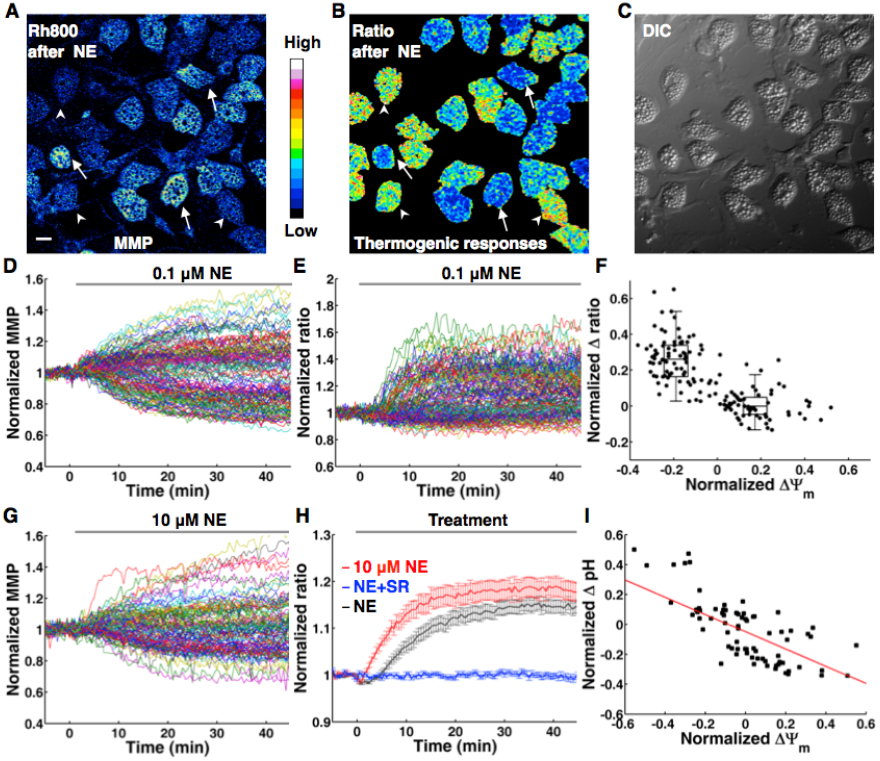
NE induces heterogeneous responses in BA. (**A-F),** NE-induced thermogenic responses in BA show negative correlation with ΔΨ_m._ **(A),** Representative MMP image of BA after 0.1 μM NE treatment. Arrowheads point to BA with mitochondrial depolarization, while arrows point to BA with mitochondrial hyperpolarization. Scale bar, 20 μm. **(B),** The corresponding thermal image (ratio of Rh800 intensity to RhB-ME intensity) after 0.1 μM NE treatment. The result shows that the low efficacy of NE-induced thermogenic responses in BA. Some BA show thermogenic responses to NE (arrow heads), while some have no responses to NE stimulation (arrows). **(C),** Differential interference contrast (DIC) image of BA. **(D)** The raw data plots of MMP show subpopulations of depolarization and hyperpolarization in 0.1 μM NE treated BA. Each colored trace represents mitochondrial membrane potential (MMP) change of single BA (n = 150). **(E)** NE-induced thermal responses in BA. Each colored trace represents a thermogenic response of single BA (n = 150). **(F)** The scatter plot of thermogenic responses (**D**) versus MMP changes (**E**) induced by NE, which shows a negative correlation (r = −0.73, n = 150, P = 2.97e-26, by Pearson’s correlation coefficient test). The box plots of the depolarization subpopulation and the hyperpolarization subpopulation show significantly different amplitudes of thermogenic responses (*P* = 1.88×10^−27^, two sided *t*-test). **(G)** The raw data plots of MMP still show two-subpopulations in 10 μM NE treated BA. Each colored trace represents a MMP change of single BA (n = 102). (**H**) Thermogenic responses in BA are evoked by 0.1 μM NE (black line, n = 150) and 10 μM NE (red line, n = 88) respectively. The blue line shows that NE-induced thermogenic response of BA is inhibited by 1 μM SR-59230A (n = 97). All data points in figures represent mean ± SEM. (**I**) A representative scatter plot of pH changes versus MMP changes induced by 0.1 μM NE, which shows a linear and negative correlation (red line, r = −0.67, n = 75, P = 3.94×10^−11^, by Pearson’s correlation coefficient test).

To test such explanation, we have checked the potential relationship between the heterogeneity of MMP changes and the extent of HSL phosphorylation at Ser563 (Figure S1), since HSL serves as a downstream lipase of β adrenergic receptors. After NE stimulation and monitoring the responses of MMP, we have retrospectively immunostained the BA with antibody against phospho-HSL and assumed that BA with high levels of phosphorylated HSL should correlate with mitochondrial depolarization. However, BA with high levels of phosphorylated HSL (arrows and red arrow head) either show mitochondrial depolarization (arrow heads) or hyperpolarization (arrows), while BA with low levels of phosphorylated HSL (white arrow heads) can show mitochondrial depolarization (Figure S1B,C and F). Secondly, treatments with NE concentration raised 100 fold to 10 μM still show two MMP populations (Figure 1G). These results suggest that the heterogeneous changes of MMP are unlikely correlated to the heterogeneity of adrenoreceptors or different activity of HSL. In addition, we have also found that all BA express UCP1 although at heterogeneous levels (Figure S2A-D), and that mitochondrial hyperpolarization is not due to the absence of UCP1 (Figure S2E-G). The different activity of HSL and heterogeneity of UCP1 may account for the diverse extent of MMP depolarization.

### Correlations among NE-induced heterogeneous responses in BA

With the thermosensitive Rhodamine B methyl ester (RhB-ME) based mito-thermotry (Xie et al., 2017), we observe that the NE-induced thermogenic responses (the fluorescent intensity ratio of Rh800 to RhB-ME) in BA indeed show negative correlation with mitochondrial electric potential difference (ΔYΨ_m_) (Figure 1D-F and movie 1-2). Our findings suggest that the heterogeneous responses of MMP in BA can index NE-induced thermogenic efficacy, and further confirm that the efficacy of NE-induced thermogenesis is low (10). Especially, even high concentration (10μM) of NE still shows low efficacy with two ΔYΨ_m_ populations (Figure 1G) and only slightly increases thermogenic responses of BA compared to the stimulation of 0.1 μM NE (Figure 1H). Meanwhile, NE-induced thermogenic responses of BA are inhibited by 1 μM SR-59230A (a potent β adrenergic receptors antagonist, Figure 1H).

Considering mitochondrial Δ Ψ_m_ is quasi-linear in physiologically relevant ranges of the pH difference (ΔpH)(21), MMP hyperpolarization might result from cytoplasmic acidification by enhanced glycolysis or NE-stimulated activities of proton pumps. Indeed, using SNARF1-AM, a ratiometric pH-indicating fluorescent probe, we reveal that the subpopulation of MMP hyperpolarization shows cytoplasmic acidification (ΔpH < 0), and ΔΨ_m_ shows negatively correlated with ΔpH (Figure 1I).

Intriguingly, the correlations among NE-induced thermogenic responses, ΔΨ_m_ and ΔpH give us a hint that mitochondrial thermogenesis of BA might be the net result of proton outflow by pumps and proton leak by UCP1.

### NE stimulation activates the proton ATPase activity of mitochondrial complex V in BA

Cytoplasmic acidification of MMP hyperpolarized BA (Figure 1I) can be not only explained by NE-acitivated proton ATPase (H^+^-ATPase) but also by enhanced glycolysis. To distinguish the difference and potential roles of the enhanced glycolysis (net ATP production) or NE-activated mitochondrial H^+^-ATPase (ATP consumption) in BA, we next examine the statuses of mitochondrial and cytoplasmic ATP concentration ([ATP]) after NE stimulation (Figure 2A-G). With FRET-based ATP indicators (mitAT1.03 and AT1.03)(22), we find that both of cytoplasmic [ATP] and majority of mitochondrial [ATP] are decreased (Figure 2C and E) after NE stimulation of BA.

**Figure 2.**
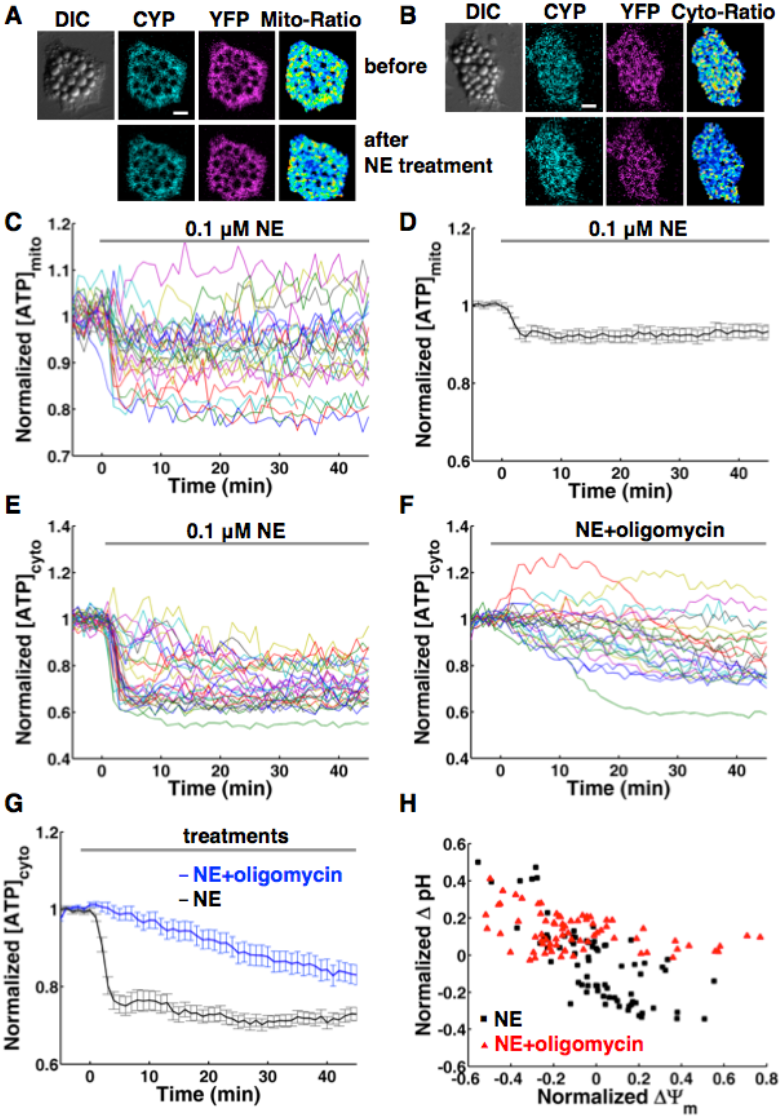
NE stimulates proton-ATPase activity of mitochondrial complex V in BA. (**A-G)** Fluorescence of AT1.03 based ratiometric measurements of mitochondrial and cytoplasmic ATP. (**A** and **B),** show fluorescence images of mitAT1.03 and AT1.03 transfected BA before and after 0.1 μM NE treatment respectively. Scale bar, 20 μm. (**C** and **D),** show raw data plots (**C**) and average plot (**D**) of mitochondrial ATP change (n = 16) in 0.1 μM NE treated BA. Each colored trace represents ATP change of single BA (**C**). (**E),** shows the raw data plots of cytoplasmic ATP (n = 29) after NE treatment. Each colored trace represents ATP change of single BA. (**F),** shows the raw data plots of cytoplasmic ATP (n = 23) after co-treatment of NE and oligomycin A. Each colored trace represents ATP change of single BA. **(G),** compares the averaged changes of cytoplasmic ATP after the treatment of NE (n = 29) or the co-treatment of NE and oligomycin A (n = 23), which show that the inhibitor (oligomycin A) of mitochondrial complex V mitigates ATP consumption induced by NE stimulation. **(H),** shows representative scatter plot of pH changes versus MMP changes induced by 0.1 μM NE with (red, n = 75) or without (black, n = 75) 10μg/mL oligomycin A pretreatment, which demonstrates that the inhibitor of mitochondrial complex V blocks the activity of proton pump. All data points in **D** and **G** represent mean ± SEM.

Clearly, these results suggest that NE stimulation activates an ATPase, which is likely a H^+^-ATPase and might play a role in the thermogenic efficacy of BA. The best candidate for H^+^-ATPase is mitochondrial ATP synthase/complex V, which can function as an ATPase to pump protons out of mitochondrial matrixes(23). ATP is not only consumed by mitochondrial ATPase but also other cellular activities. Therefore, it is necessary to further check whether complex V is the ATPase activated by NE. Thus, we use oligomycin A, a specific inhibitor of complex V, and reveal that oligomycin A mitigates ATP consumption induced by NE stimulation (Figure 2F and G). These results confirm that NE activates the ATPase function of mitochondrial complex V.

The role of mitochondrial complex V as ATPase is quite provocative. For example, where does ATP come from? The source of ATP production can be provided by NE-stimulated metabolism of glucose, since it has been reported and well known that glucose uptake is increased in BA or BAT by one or two orders of magnitude in human (12 folds) and rat (110 times) under cold stimulation(24),(25). NE-stimulated metabolism in BA is indeed supported by accelerated-metabolic status represented with homogenous redox changes (Figure S3A-H), which are endogenous autofluorescences and ratio measurement of FAD and NADH (16). In addition, interestingly illustrated in Figure 2F, a few of BA pretreated with oligomycin A show transiently increased cytoplasmic [ATP], which support accelerated cytosol metabolism for ATP production after NE stimulation.

In addition, to further check whether the ATPase function of complex V causes cytoplasmic acidification after NE stimulation, we use oligomycin A to inhibit complex V again and reveal that almost all of BA show cytoplasmic alkalizations (ΔpH > 0) and that majority of BA show MMP depolarization after NE stimulation (Figure 2H). Thus, these findings further demonstrate and confirm that NE stimulation activates the H^+^-ATPase function of complex V rather than glycolysis for cytoplasmic acidification. Together, our findings suggest that the H^+^-ATPase activity of complex V is a key factor to antagonize proton leak by UCP1, impact on the net cytoplasmic ΔpH (Figure 1I and Figure 2H) and modulate the efficacy of NE-induced BA thermogenesis.

### ATP as a sympathetic cotransmitter can increase the NE-induced thermogenic efficacy

Our observations suggest that more factors other than NE alone are needed for the high efficacy of evoking thermogenesis in BAT. Considering ATP is also a neurotransmitter and co-released with NE when sympathetic nerve firing is evoked(26), we examine whether ATP is a co-factor needed for NE to efficiently evoke thermogenesis of BA (Figure 3). As illustrated in Figure 3, majority of BA show thermogenic responses (Figure 3A-C) to NE and ATP co-stimulation (movie 3-4). The co-treatment with NE and ATP also increases the percentage (in mean ± SD) of MMP depolarization subpopulation (72.8% ± 13.3%) in BA compared to NE treatment alone (55.6% ± 16.1%, P = 0.0514, one sided t-test) (Figure 3D-E). NE and ATP co-stimulation markedly increases the amplitudes of thermogenic responses (Figure 3F) in BA compared to NE treatment alone (Movie 1-4). Whereas, BA only show slightly and relative flat responses to ATP treatment alone (Figure 3F).

**Figure 3.**
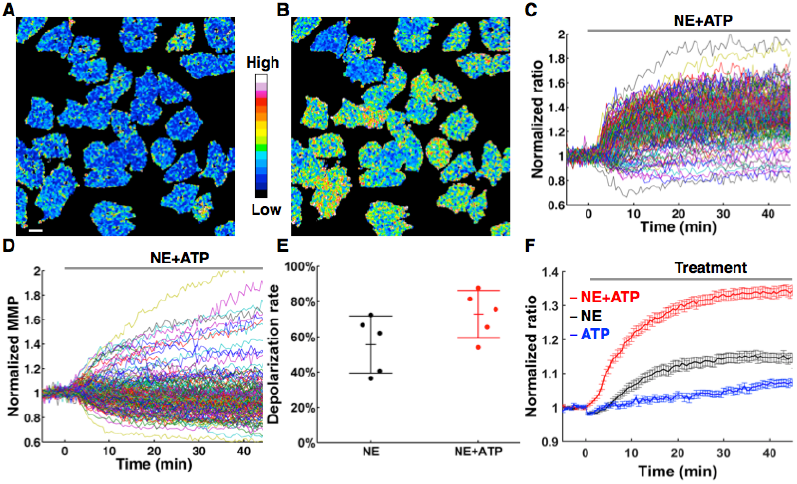
Sympathetic cotransmitter ATP enhances NE-induced thermogenic efficacy in BA. (**A and B**), Representative thermal images of BA before and after 0.1 μM NE and 10 μM ATP co-treatment respectively. Scale bar, 20 μm. **(C),** 0.1 μM NE and 10 μM ATP induced thermal responses in BA. Each colored trace represents a thermogenic response of single BA (n= 158). **(D),** The dynamic plots of MMP demonstrate that majority of BA show MMP depolarization after NE and ATP co-treatment. Each colored trace represents a MMP dynamic of single BA (n = 158). **(E),** shows scatter plots of MMP depolarization percentage in 0.1 μM NE without (black) or with (red) ATP treated experiments. NE and ATP co-treatment increases the percentage of MMP depolarization (72.8% ± 13.3%) compared to NE treatment alone (55.6% ± 16.1%). Error bars in figure represent mean ± SD. **(F),** 0.1 μM NE without (black line, also in Fig. 1H) or with 10 μM ATP (red line, n = 158) induced thermogenesis in BA. The blue line shows the control results of ATP treatment alone in BA (n = 67). All data points in **F** represent mean ± SEM.

To elucidate the enhancement effect of sympathetic cotransmitter ATP via purinergic receptors, we check the statuses of cytoplasmic and mitochondrial Ca^2+^ concentration ([Ca^2+^]) with ratiometric probe Fura2-AM and indicator 4mtD3cpv(27) respectively (Figure 4 and S4). We find that extracellular ATP evokes transient elevation of cytoplasmic [Ca^2+^] (Figure 4A and C) in BA through multiple P2 receptors (Figure S4 and Movie 3) in line with previous reports(28). Whereas, we find that extracellular ATP evokes transient peaked and steady elevation of mitochondrial [Ca^2+^] (Figure 4B and D), which can upregulate mitochondrial oxidative phosphorylation(29).

**Figure 4.**
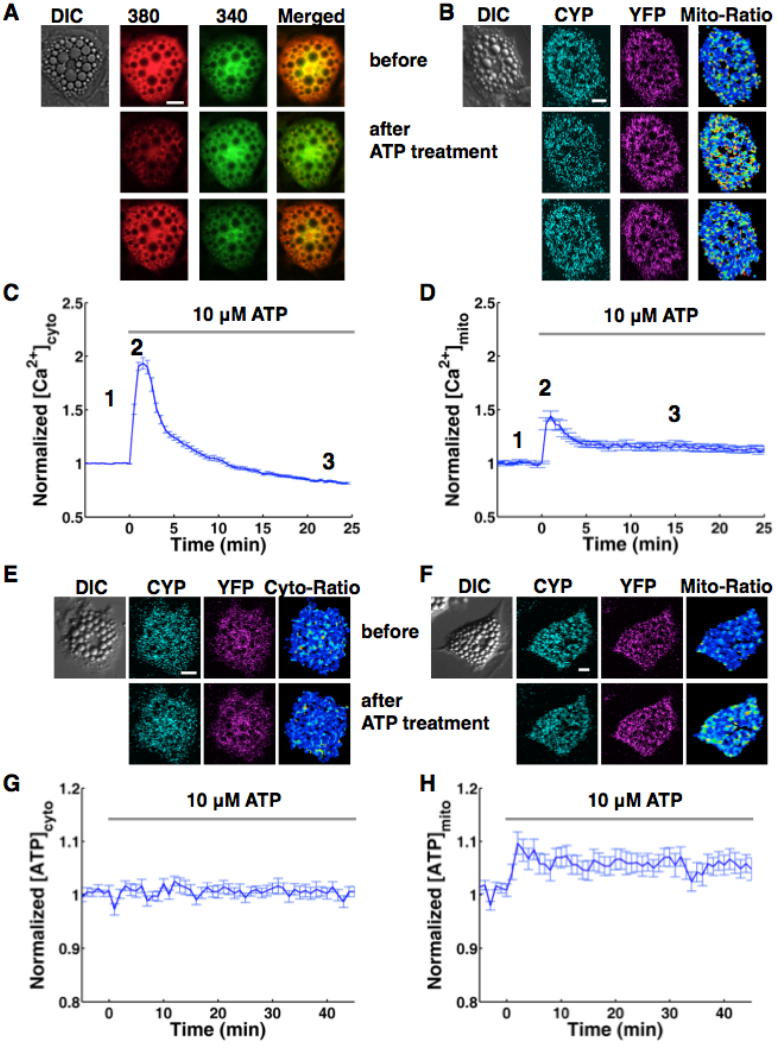
Sympathetic cotransmitter ATP enhances NE-induced thermogenic efficacy in BA through modulating the function of mitochondrial complex V. (**A-D),** Sympathetic cotransmitter ATP increases cytoplasmic and mitochondrial [Ca^2+^] in BA, which are both fluorescence-based ratiometric measurements. Scale bars, 20 μm. **(A),** shows fluorescence images (excited by 380 nm or 340 nm laser) of Fura2-AM stained BA before and after 10 μM ATP treatment respectively, and their positions in panel **C** are labeled with 1, 2 and 3 respectively. **(B),** shows fluorescence images of 4mtD3cpv transfected BA before and after 10μM ATP treatment respectively, and their positions in panel **D** are labeled with 1, 2 and 3 respectively. (**C** and **D),** show that ATP induces cytoplasmic (n = 63) and mitochondrial (n = 14) [Ca^2+^] transient in BA respectively. (**E-H),** Fluorescence of AT1.03 based ratiometric measurements of cytoplasmic and mitochondrial ATP concentration induced by extracellular ATP. (**E** and **F),** show fluorescence images of AT1.03 and mitAT1.03 transfected BA before and after 10 μM ATP treatment respectively. Scale bars, 20 μm. (**G** and **H)** show the averaged change of cytoplasmic (n = 28) and mitochondrial (n = 31) ATP by 10 μM ATP treatment. All data points in **C**, **D**, **G** and **H** represent mean ± SEM.

Therefore, we examine the states of cytoplasmic and mitochondrial [ATP] after ATP stimulation (Figure 4E-H). We observe a steady elevation of mitochondrial [ATP] while no apparent change of cytoplasmic [ATP] after extracellular ATP stimulation (Figure 4E-H). These findings suggest that sympathetic co-transmitter ATP increases the ATP synthase activity of complex V for ATP production (Figure 4H) by elevating mitochondrial [Ca^2+^] (Figure 4D).Importantly, the thermogenic enhancing function (Figure 3) of ATP as sympathetic co-transmitter of NE can explain the discrepancy of cold and sympathomimetics activities on human BAT *in vivo*(30).

## DISCUSSION

### Mitochondrial complex V is a deterministic factor in NE-induced thermogenic efficacy

How does the H^+^-ATPase activity of complex V modulate the thermogenic efficacy of BA? A first straightforward explanation is endothermic characteristic feature of ATP hydrolysis by mitochondrial complex V(31), which pumps dissipated proton back and restore energy in mitochondrial proton gradient. In Figure 1E, there are some interesting and supporting results, where very a few BA show endothermic profiles rather than exothermic profiles to NE stimulation. The endothermic profiles could be a stimulus artifact but unlikely. Secondly, H^+^-ATPase activity of complex V results in cytoplasmic acidification (Figure 1I and Figure 2H), which can enhance the binding and inhibition of purine nucleotides to UCP1(32).

Why need the function of complex V as H^+^-ATPase by NE stimulation? The physiological role of the H^+^-ATPase function is likely to avoid unnecessary and undesired heat production since NE is either secreted by adrenal medullae or released by sympathetic nervous system, which can be also excited by mental or physical stress (33). Accordingly, for cold stimulation, our observations suggest that at least two factors (NE and ATP) are needed for the high efficacy and specificity of evoking thermogenesis in BAT.

In this study, we have observed that NE evokes both homogenous responses (of redox ratio, cytosol ATP consumption and calcium dynamics) and heterogeneous changes (in MMP, cytosol pH and thermogenesis). Consequently, we have demonstrated that the heterogeneous changes and thermogenic efficacy are modulated by the functional status of mitochondrial complex V. Our findings suggest that mitochondrial malfunctions or impairments in sympathetic system or BAT can be pathologic reasons for inefficient energy expenditure and developing obesity. Additionally, our current observations also open new doors and pave ways to reconsider targets and strategies of fully utilizing brown adipocytes thermogenic capacity for the treatments of obesity, diabetes and abnormal lipid metabolism.

## MATERIALS and METHODS

Data are pooled from or repeated for each condition at least three independent experiments. No statistical methods were used to predetermine sample size. All data points in figures represent mean ± SEM except Figure 3E, which data in text or figures represent mean ± SD. More detailed information of the Experimental Procedures can be found in the Supplemental Data.

### Isolation and primary culture of brown adipocytes

Brown adipocytes (BA) were isolated from 3-4weeks male C57BL/6J mice, with a procedure similar to that described by Lucero *et al.* (13). Briefly, mice were kept at 4 °C for overnight with free access to food and water to deplete stored lipid in brown adipose tissues (BAT). The mice were then sacrificed by cervical dislocation and swabbed with 75% ethanol. Interscapular BAT was isolated and placed in isolation buffer (DMEM with 4% NCS). The tissue was minced and digested with 0.2% collagenase type II in a shaking water bath at 37 °C for 30 min (14). After digesting, discarded the reaction mixture, and the tissue was washed with isolation buffer. Dissociated cells by gently triturating with fire-polish pipettes and washed by centrifugation in PBS. After the final washing, the cells were plated onto 12 mm coverslips (∼3×10^4^ cells per coverslip). Coverslips were precoated with matrigel. After 2 hours, 2 ml plating medium (DMEM supplemented with 5% FBS, 100 units/ml penicillin and 100 μg/ml streptomycin) was added to each 35 mm dish. From the second day in culture, half of the medium was replaced with feeding medium (plating medium supplemented with Ara-C to inhibit fibroblast proliferation, 2 μM final concentration of Ara-C) every 2 days. Cells were maintained at 37 °C in a humidified atmosphere of 95% air and 5% CO_2_, and used for imaging at 3ߨ days *in vitro*.

### Cell transfection

BA were washed with PBS and resuspended in electroporation solution (20 mM Hepes, 135 mM KCl, 2 mM MgCl_2_, 0.5% Ficoll 400, 1% DMSO, 2 mM ATP and 5 mM glutathione, pH 7.6) with 20 μg of plasmid DNA (15). Two pulses (115 V, 10 ms duration) with 1s interval were delivered to electroporate cells (1-1.5×10^6^ cells/ml) with an ECM 830 Square Wave Electroporation System (Harvard Apparatus, Inc., USA). Then, cells were seeded onto matrigel-coated coverslips. BA were maintained in DMEM supplemented with 5% FBS until imaging.

### Thermogenic study

All imagings were performed using confocal microscope with a 40×/0.95 objective (Olympus) for time-lapse imaging and a 100×/1.4O objective (Olympus) for high-resolution imaging (Xie et al., 2017). Cells were co-stained with RhB-ME and Rh800 (20 nM dyes for time-lapse imaging, 50 nM dyes for high resolution imaging) in Tyrode’s solution (in mM: 10 Hepes, 10 glucose, 3 KCl, 145 NaCl, 1.2 CaCl_2_, 1.2 MgCl_2_, pH 7.4) for 1 h at 33 °C. The pseudocolor of RhB-ME channel is red (excited at 559 nm and collected at 575-620 nm), and Rh800 channel is green (excited at 635 nm and collected at 655-755 nm). All images were collected at 512 × 512 (for time-lapse imaging) and 1600 ×1600 (for high resolution imaging) pixels resolution (12 bit). All data analysis was performed with MATLAB (MathWorks Inc. USA) or ImageJ (NIH, USA).

### Cytoplasmic pH study

Cytoplasmic pH imagings were performed using confocal microscope with a 40×/0.95 objective (Olympus) for time-lapse imaging. Cells were co-stained with 5μM SNARF1-AM and 20 nM Rh800 in Tyrode’s solution for 30 min at 37 °C. Then wash out SNARF1-AM using Tyrode’s solution (with 20 nM Rh800). The pseudo color of acidification channel is red (excited by 561 nm laser and collected with 575-600 nm band pass filter), and alkalization channel is green (simultaneously collected with 600-675 nm band pass filter); Rh800 channel is magenta (sequentially excited by 635 nm laser and collected with 655-755 nm band pass filter). Ratiometric values of acidification channel to alkalization channel were calculated pixel by pixel to represent the relative pH value of the sample. The other conditions of imaging, data acquisition and analysis for cytoplasmic pH study were the same as the conditions in thermogenic study.

### Redox (FAD/FAD+NADH) measurement

Redox (FAD/FAD+NADH) imagings were performed using a customized fluorescence microscope with a 40×/0.8W objective (Olympus) for time-lapse imaging. The endogenous autofluorescence images(16) were excited with Optoscan monochromator (Cairn Research Ltd., UK). The pseudocolor of FAD channel is red (excited by 430 nm light with 20 nm bandwith and collected with 525-575 nm band pass filter), while NADH+FAD channel is green (sequentially excited by 340 nm light with 20 nm bandwith and collected with 420 nm long pass filter). And the fluorescence images were acquired with Evolve 512 EMCCD (Photometrics Ltd., UK).

Time-lapse imagings of BA were performed in 4 mL Tyrode’s solution at 33 °C. 60 frames were recorded in a time-lapse imaging at 30 seconds interval. NE (0.1 μM) was injected as soon as the 11th frame of the time-lapse imaging for Redox studies in BA. The other conditions of imaging, data acquisition and analysis for Redox study were the same as the conditions in thermogenic study.

### Cytoplasmic [Ca^2+^] study

Cytoplasmic [Ca^2+^] imagings were performed using the customized fluorescence microscope with a 40×/0.8W objective (Olympus) for time-lapse imaging and Optoscan monochromator (Cairn Research Ltd., UK) as light source. Cells were stained with 5μM Fura2-AM in Tyrode’s solution for 30 min at 37 °C. Then wash out Fura2-AM using Tyrode’s solution. The pseudocolor of 380 nm channel is red (excited by 380 nm light with 10 nm bandwith and collected with 505-535 nm band pass filter), and 340 nm channel is green (sequentially excited by 340 nm light with 10 nm bandwith and also collected with 505-535 nm band pass filter). Ratiometric values of 340 nm channel to 380 nm channel were calculated pixel by pixel to represent the relative [Ca^2+^] of the sample. The other conditions of imaging, data acquisition and analysis for cytoplasmic [Ca^2+^] study were the same as the conditions in Redox study.

### Mitochondrial [Ca^2+^] study

Mitochondrial [Ca^2+^] imagings were performed using confocal microscope with a 40×/0.95 objective (Olympus) for time-lapse imaging. Cells were transfected with 4mtD3cpv (from Dr. R. Tsien) using ECM 830 Square Wave Electroporation System (Harvard Apparatus, Inc., USA). The pseudocolor of CFP channel is cyan (excited by 440 nm laser and collected with 480-495 nm band filter), and YFP channel is magenta (simultaneously collected with 505-605 nm band filter). Ratiometric values of YFP channel to CFP channel were calculated pixel by pixel to represent the relative [Ca^2+^] of the sample. The other conditions of imaging, data acquisition and analysis for mitochondrial [Ca^2+^] study were the same as the conditions in cytoplasmic [Ca^2+^] study.

### Cytoplasmic and mitochondrial [ATP] study

[ATP] imagings were performed using confocal microscope with a 40×/0.95 objective (Olympus) for time-lapse imaging. Cells were transfected with AT1.03 for cytoplasmic [ATP] or mitAT1.03 for mitochondrial [ATP] (from Dr. H. Noji and Dr. H. Imamura) using ECM 830 Square Wave Electroporation System (Harvard Apparatus, Inc., USA). The pseudocolor of CFP channel is cyan (excited at 440 nm and collected at 480-495 nm), and YFP channel is magenta (simultaneously excited at 440 nm and collected at 505-605 nm). Ratiometric values of YFP channel to CFP channel were calculated pixel by pixel to represent the relative [ATP] of the sample. The other conditions of imaging, data acquisition and analysis for [ATP] study were the same as the conditions in thermogenic study.

## Supplemental Data

Supplemental Data include Supplemental four figures and five movies.

## Author Contributions

T.R.X. and C.F.L. designed and conducted the experiments and manuscript writing. T.R.X. did live cell imaging and data analysis. C.F.L. did primary cell culture and imaging. J.S.K. built the custom microscopy, developed the idea, directed the study and wrote the paper. All authors participated in discussions.

## Acknowledgements

We thank the following people for their help: Dr. H. Noji and Dr. H. Imamura for AT1.03 and mitAT1.03 plasmids; Dr. R. Tsien for 4mtD3cpv plasmid.

## Supporting online material

**Figure S1.**
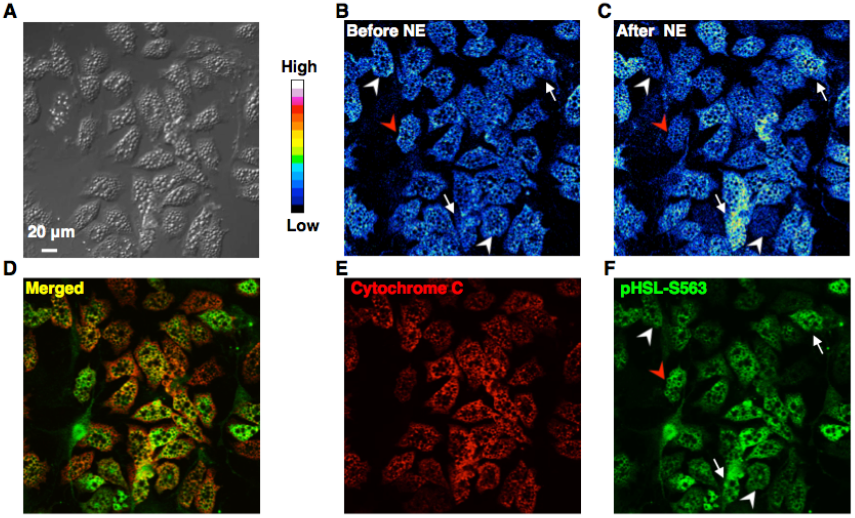
NE-induced heterogeneous changes of mitochondrial membrane potentials (MMP) are not correlated to the heterogeneity of phospho-HSL levels. (**A-C),** Merged confocal images of mitochondria in BA stained with 20 nM mitochondrial marker Rh800. Scale bar, 20 μm. **(A),** Differential interference contrast (DIC) image of multilocular BA. (**B** and **C),** Representative images of mitochondrial membrane potential (represented with Rh800 intensity) are shown for the moments before and after 0.1 μM NE treatment respectively. Arrow heads point to BA with mitochondrial depolarization, while arrows point to BA with mitochondrial hyperpolarization. (**D-F),** Heterogeneous changes of MMP in BA are not correlated with phosphorylated levels of HSL. Retrospectively immunostained images of BA after NE treatment. **(D),** Merged image of BA stained with antibodies against cytochrome C (red, **E**) and phosphorylated HSL (green, **F**). Red arrow head points to BA with mitochondrial depolarization (**B** and **C**) and high level of phosphorylated HSL (**F**), while white arrows point to BA with mitochondrial hyperpolarization (**B** and **C**) and high level of phosphorylated HSL (**F**). Meanwhile, arrow heads show BA with mitochondrial depolarization (**C**) and low level of phosphorylated HSL (**F**).

**Figure S2.**
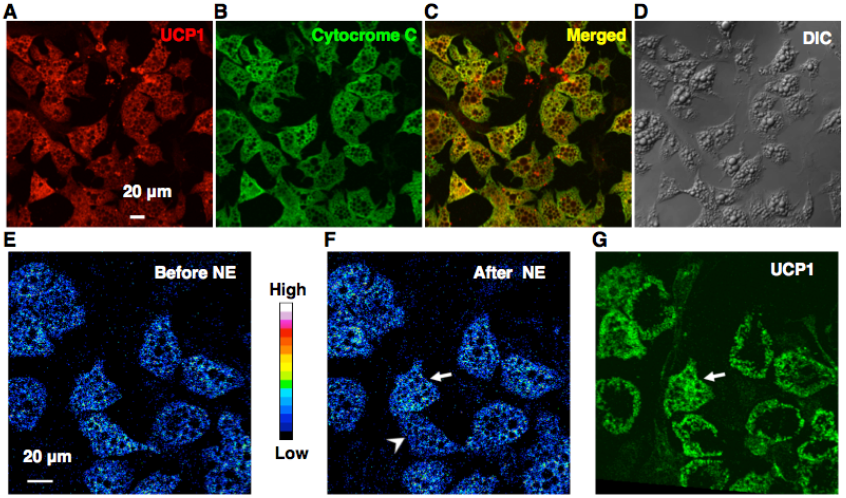
Hyperpolarization of MMP in BA is not due to the absence or heterogeneity of UCP1 levels. (**A-D),** Confocal images of BA immunostained with antibodies against UCP1 (red, **A**) and Cytochrome C (green, **B**). Scale bar, 20 μm. **C,** Merged image of immunostained BA. (**D),** Differential interference contrast (DIC) image of BA. (**E** and **F),** Hyperpolarization of MMP in BA is not correlated with expression levels of UCP1. Representative images of mitochondrial membrane potential (represented with Rh800 intensity) are shown for the moments before and after 0.1 μM NE treatment respectively. (**G),** Confocal images of BA retrospectively immunostained with antibody against UCP1 after NE treatment (**E** and **F**). Arrow points to BA with mitochondrial hyperpolarization with high level of UCP1, while arrow head shows BA with mitochondrial depolarization.

**Figure S3.**
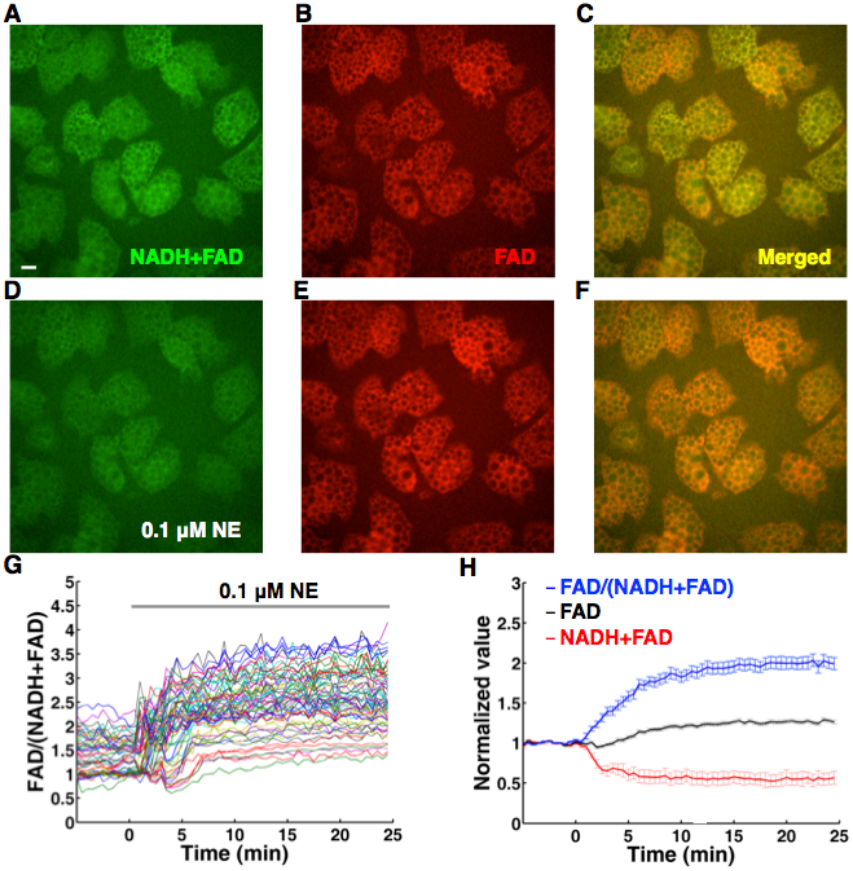
Redox ratio measurement for NE stimulation in BA. (**A-H),** Autofluorescence of FAD and NADH based redox ratio measurements. Scale bar, 20 μm. (**A** and **D),** show autofluorescence images of FAD + NADH signals before and after 0.1 μM NE treatment respectively, while (**B** and **E)** show autofluorescence images of FAD before and after NE treatment respectively. (**C** and **F)** are merged redox ratio images of FAD (red) to FAD+NADH (green). (**G),** The raw data plots of redox ratio change in 0.1 μM NE treated BA. Each colored trace represents redox change of single BA (n =60). **(H)** shows the normalized averaged FAD, FAD+NADH and redox change after NE treatment. All data points in (**H)** represent mean ± SEM.

**Figure S4.**
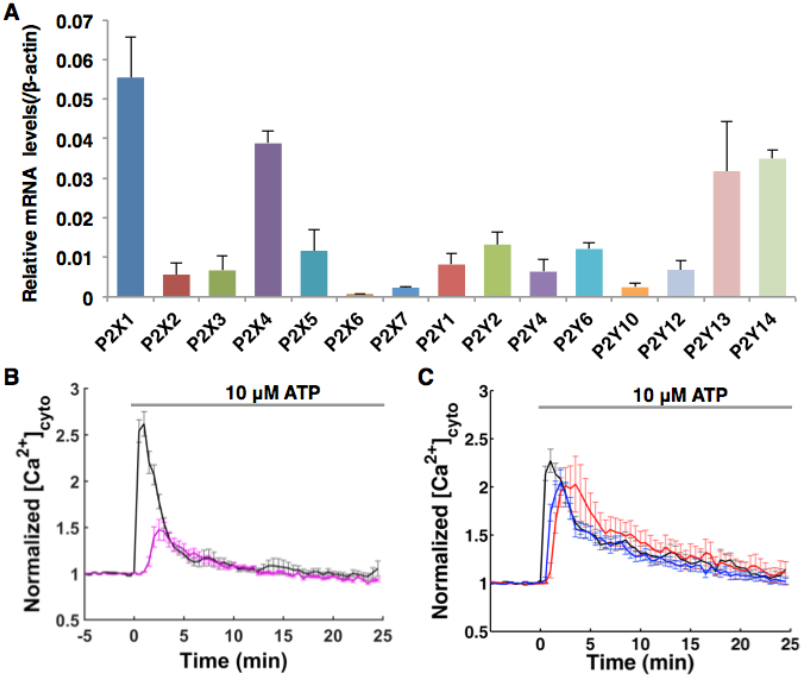
Sympathetic cotransmitter ATP increases intracellular [Ca^2+^] in BA through multiple P2 receptors. (**A**), The quantitative-PCR results show the expression levels of P2 receptors in BA, including P2X1, P2X2, P2X3, P2X4, P2X5, P2X6, P2X7, P2Y1, P2Y2, P2Y4, P2Y6, P2Y10, P2Y12, P2Y13 and P2Y14. (**B** and **C),** show that ATP induces cytoplasmic [Ca^2+^] transient in BA can be dramatically decreased with multiple inhibitors of P2 receptors **(B)**, and less affected by single inhibitor of P2X receptor (**C**). **(B)** shows that the combination of multiple inhibitors (10 μM PPADS, a nonselective inhibitor of P2X receptor; 2.5 μM PSB 0739, an antagonist of P2Y12; 0.5 μM MRS 2279, an inhibitor of P2Y1; 100 μM Suramin, a nonselective inhibitor of P2Y receptor) can largely decrease the cytoplasmic [Ca^2+^] transient induced by ATP stimulation (Control, black, n = 25; multiple inhibitors, magenta, n = 21). **(C)** shows that single inhibitor has little effect to reduce cytoplasmic [Ca^2+^] transient induced by ATP stimulation (Control, black, n= 17), such as a selective antagonist of P2X1 (10 μM PPNDS, blue, n = 14) or nonselective inhibitor of P2X receptor (10 μM PPADS, red, n = 13). All data points in **A-C** represent mean ± SEM.

### 3. Supplemental Video Legends

#### Supplemental Video 1 | Subpopulations of NE-induced thermogenesis in BA

The ratiometric pseudocolor movie of 0.1μM NE-induced thermal responses in BA. Scale bar, 20 μm.

#### Supplemental Video 2 | NE-induced heterogeneous responses in BA

The raw channel data of supplementary video 1 show BA subpopulations after NE treatment. Red represents the channel of thermosensitive RhB-ME, while green represents thermoneutrol Rh800. Scale bar, 20 μm.

#### Supplemental Video 3 | NE and ATP co-induced thermogenesis in BA

The ratiometric pseudocolor movie of co-treatment with 0.1 μM NE and 10 μM ATP-induced thermal responses in BA. Scale bar, 20 μm.

#### Supplemental Video 4 | Raw data of NE and ATP co-induced thermogenesis in BA

The raw channel data of supplementary video 3 show BA subpopulations after NE and ATP co-treatment. Red represents the channel of thermosensitive RhB-ME, while green represents thermoneutrol Rh800. Scale bar, 20 μm.

#### Supplemental Video 5 | ATP-induced cytoplasmic [Ca^2+^] transient in BA

The merged pseudocolor movie of 10 μM ATP-induced cytoplasmic [Ca^2+^] transient in BA. Scale bar, 20 μm.

### 4. Supplemental Materials

#### General materials

Rhodamine B (RhB), Rhodamine 800 (Rh800), and carbonyl cyanide m-chlorophenyl hydrazone (CCCP), rotenone, oligomycin A, adenosine triphosphate (ATP), anti-UCP1 (U6382, rabbit), collagenase type II, cytosine arabinoside (Ara-C), Ficoll 400, glutathione, glucose, Hepes, KCl, NaCl, MgCl_2_ and CaCl_2_ were purchased from Sigma-Aldrich Corporation (USA). Norepinephrine (NE) and Dimethyl Sulfoxide (DMSO) were purchased from Santa Cruz Biotechnology, Inc. (USA).SR-59230A was purchased from Abcam Inc. (UK). SNARF1-AM, Fura2-AM, Dulbecco’s Modified Eagle Medium (DMEM), fetal bovine serum (FBS), newborn calf serum (NCS), ALEXA-488 goat anti-rabbit, ALEXA-555 goat anti-mouse Phosphate-Buffered Salines, pH7.4 (PBS) and Penicillin Streptomycin (Pen Strep) were purchased from Thermo Fisher Scientific Co., Ltd. (USA). Anti-Phospho-HSL (Ser563, 4139, rabbit) was purchased from Cell Signaling Technology (USA). Matrigel and anti-cytocrome C (556432, mouse) were purchased from BD Biosciences Company (USA). Coverslips were purchased from Glaswarenfabrik Karl Hecht GmbH & Co KG (Germany).

C57BL/6J mice were purchased from Sino-British SIPPR/B&K Lab Animal Ltd., Shanghai (China). All experimental procedures and protocols were approved by the Institutional Animal Care and Use Committee of the Institute for Nutritional Sciences, Shanghai Institutes for Biological Sciences, Chinese Academy of Science.

All confocal imaging was performed using confocal microscope FV1000 (Olympus Corporation, Japan). The other fluorescence imagings were performed using a customized fluorescence microscope (BX61WI, Olympus Corporation, Japan) equipped with Optoscan monochromator (Cairn Research Ltd., UK), customized emission filters (Semrock, USA) and Evolve 512 EMCCD (Photometrics Ltd., USA), which were all controlled with customized Micro-Manager software.

#### Thermogenic study

All imagings were performed using confocal microscope with a 40×/0.95 objective (Olympus) for time-lapse imaging and a 100×/1.4O objective (Olympus) for high-resolution imaging. Cells were co-stained with RhB-ME and Rh800 (20 nM dyes for time-lapse imaging, 50 nM dyes for high resolution imaging) in Tyrode’s solution (in mM: 10 Hepes, 10 glucose, 3 KCl, 145 NaCl, 1.2 CaCl_2_, 1.2 MgCl_2_, pH 7.4) for 1 h at 33 °C (10). The pseudocolor of RhB-ME channel is red (excited at 559 nm and collected at 575-620 nm), and Rh800 channel is green (excited at 635 nm and collected at 655-755 nm). All images were collected at 512 × 512 (for time-lapse imaging) and 1600 ×1600 (for high resolution imaging) pixels resolution (12 bit).

To minimize the heat influence by perfusion solution, time-lapse imaging of BA was performed in 2 mL Tyrode’s solution rather than in perfusion system. To minimize bleaching and damage to live BA in time-lapse imaging, the lowest intensity of lasers with largest pinhole setting and the shortest scanning time were used, and 101 frames were recorded in a time-lapse imaging at 30 seconds interval. NE (0.1 μM), ATP (10 μM), or vehicle was injected as soon as the 11th frame of the time-lapse imaging for thermogenesis studies in BA. SR-59230A (1 μM), or oligomycin (10 μg/mL) was injected 15 min before the time-lapse imaging in pharmacological tests.

After background being removed, ratiometric values of Rh800 channel to RhB-ME channel (simultaneously excited by 635 nm and 559 nm lasers) were calculated pixel by pixel to represent the thermal response of the sample. For noise reduction, the pixels with signal-to-noise ratios less than 1.5 were excluded and 5.5 moving average was used before ratiometric process. According the three-sigma rule, the outlier (99.7% tolerance interval) of the ratios was also excluded. The ratio of each cell was the average ratio of all pixels representing the cell. Since we focus on the thermal responses, the mean ratio of steady state before drug treatments was used to normalize every data point for each cell. All data analysis was performed with MATLAB (MathWorks Inc. USA) and ImageJ (NIH, USA).

#### Primers for quantitative-PCR analysis of P2 receptors in BA

Total RNA of cultured brown adipocytes were extracted and prepared with Trizol (Thermo Fisher, Cat No: 15596-026). First strand cDNAs were synthesised using oligo(dT)18 primers (Thermo Fisher, First Strand cDNA Synthesis Kit, Cat No: K1612) and RT-PCR reagent, which contains 4.6 μl cDNA (100 ng/μl), 0.2 μl each primer (10 μM) and 5 μl SYBR green mix (Thermo Fisher, Cat No: 4367659). The paired primers of *P2x* and *P2y* were listed as below:

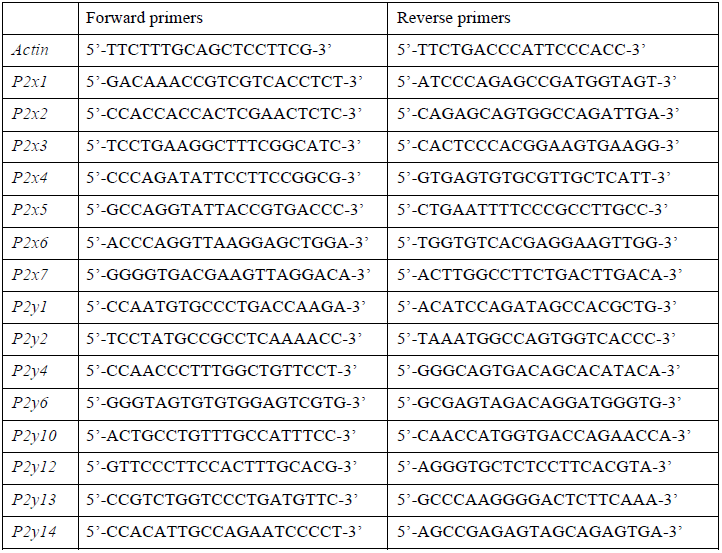

All the reagents were pipetted into 384-well plate, gently spun down, and placed in 7500 Real-Time PCR System (ABI). Samples were Incubated at 95°C for 10 min, followed by 40 cycles of 95°C for 25 s, 60°C for 30 s, and 72°C for 30 s. Data were collected and analyzed by being normalized with actin.

